# Detection and measurement of butterfly eyespot and spot patterns using convolutional neural networks

**DOI:** 10.1101/2022.03.21.485083

**Authors:** Carolina Cunha, Antónia Monteiro, Margarida Silveira

## Abstract

Butterflies are increasingly becoming model insects where basic questions surrounding the diversity of their color patterns are being investigated. Some of these color patterns consist of simple spots and eyespots. To accelerate the pace of research surrounding these discrete and circular pattern elements we developed two distinct convolutional neural networks (CNNs) for detection and measurement of butterfly spots and eyespots on digital images of butterfly wings. We tested the accuracy of the detection and of the area measurements using manual identifications and measurements. These methods were able to identify and distinguish marginal eyespots from spots, as well as distinguish these patterns from less symmetrical patches of color. In addition, the measurements of an eyespot’s central area and surrounding rings were highly accurate. These CNNs offer improvements of eyespot/spot detection and measurements relative to previous methods because it is not necessary to mathematically define the feature of interest. All that is needed is to point out the images that have those features to train the CNN.

**Author summary:** We developed two distinct convolutional neural networks (CNNs) for detection and measurement of butterfly spots and eyespots on digital images of butterfly wings. We tested the accuracy of the detection and of the area measurements using manual identifications and measurements. Our methods were able to identify and distinguish marginal eyespots from spots, as well as distinguish these patterns from less symmetrical patches of color. In addition, the measurements of an eyespot’s central area and surrounding rings were highly accurate.

## Introduction

Eyespots are salient color pattern stimuli with multiple rings of contrasting colors that mimic vertebrate eyes. They are used by multiple animals primarily to intimidate or startle predators or to deflect predator attacks to dispensable areas of the body [1]. Eyespots have been studied primarily in the lepidoptera, where different modes of defense are found in different species, and where eyespots are also used in sexual signaling [2–4]. Eyespots in nymphalid butterflies have a single origin [5], and they may have evolved from simpler pattern elements, spots [6]. Spots are simple patches of color contrasting against the background color of the wing. It is unclear how many times spots have evolved independently in butterflies and what exact ecological function they serve.

The accurate measurement of spots and eyespots has become a routine task for researchers who study the ecological role of these traits in butterflies. For example, different sizes of eyespots found in males and females, hinted at their role in sexual signaling [7], whereas different sizes of eyespots in dry and wet seasons forms of the same species of butterfly hinted at the presence of different predator guilds in each season shaping eyespot size [8].

Measuring spots and eyespots has also become important for researchers who explore mechanistic questions about their development. For instance, in order to probe the mechanism of sexual size dimorphism or seasonal phenotypic plasticity, researchers use a variety of hormone and drug injections during development to test how they affect the final size of the eyespots [9, 10]. Eyespot and spot measurements are also performed to test the role of candidate genes in the development of these traits using genetic perturbations [11–13]. Having the ability to perform quick measurements on spots/eyespots is, thus, useful to accelerate the pace of research around these traits.

Previous approaches for automatic eyespot detection and measurement [14] relied on a sliding window approach where carefully selected features exploiting symmetry and circularity were measured. These features were then fed to an SVM classifier for detection, after which the different circular rings were measured with a 1D Hough Transform for circle detection.

In this work, we use convolutional neural networks (CNN) [15] for both spot/eyespot detection and measurements. CNNs are currently the most widely used deep learning algorithms for image analysis, having outperformed traditional algorithms in many image analysis problems like image classification, object detection or segmentation [16, 17]. While CNNs have been used in the past for purposes of butterfly species identification [18–22], here we use them to identify specific wing patterns, regardless of species identity. The main advantage of CNNs over previous methods is their ability to automatically extract from the images the most relevant features of the patterns of our choice, in this case spots and eyespots, requiring no manual feature extraction. We first use a dataset of images of different species of butterflies with a variety of spots and eyespots for training and testing purposes. Then we test our CNN to detect eyespots in a different image dataset consisting of images of many individuals of a single species. In this second image dataset we also train and test a new CNN to segment and measure different eyespot areas.

## Materials and methods

### Data

Two different datasets were used in our experiments. For the spot and eyespot detection task we used a dataset (dataset1) with 4707 butterfly images of different species, 690 of which were images of butterflies without any spots or eyespots, and the remaining 4017 contained one or more spots or eyespots. For each butterfly image containing spots/eyespots, the right-wing pattern elements had been identified manually in two previous studies [5, 6] and their center position and size recorded. The type of pattern element was also identified. Fig 1 illustrates the variability of the different types of pattern elements. Table 1 reports the number of pattern elements of each type that is present in dataset1.

**Table 1.** Number of pattern elements from each type in dataset1 (see illustrations and definitions of pattern elements in Fig 1).

| Spots | Eyespots |
| --- | --- |
| 17 382 | 16571 |

**Fig 1.**
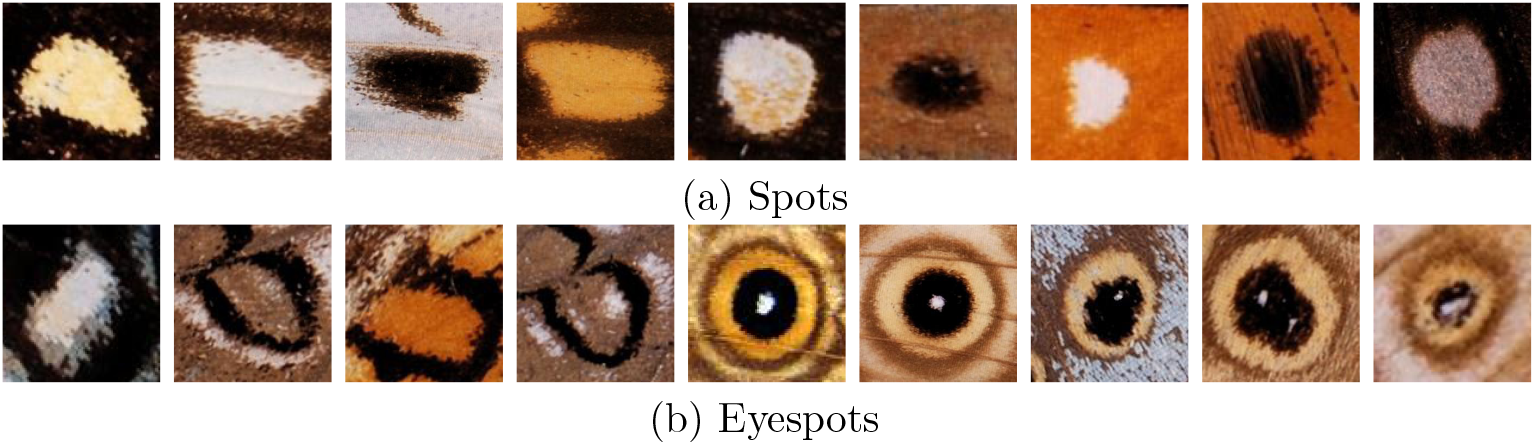
Examples of the types of pattern element present in dataset 1. Spots are pattern elements that develop a single spot of color. Eyespots are pattern elements that develop spots and rings of color. These include discal and marginal eyespots. The first are eyespots that develop around a cross-vein and are found in the center of the wing, and the second develop closer to the margin of the wing.

For the eyespot measurement task we used a different dataset (dataset 2) containing 64 images of the ventral forewing of a single species, *Bicyclus anynana*. In each image, two marginal eyespots, a small and a bigger one, had been identified manually and their center coordinates recorded. Additionally, for each eyespot, the total area, up to the outer perimeter of the orange ring, and the white center area had been measured in *mm*^2^.

### Image Analysis

The analysis comprised two steps that use CNN’ s: (1) spot/eyespot detection and (2) spot/eyespot measurements. First, the butterfly image is fed to the detection CNN, which provides as output the bounding boxes of the detected spots/eyespots and corresponding types. Then, for each pattern element that was detected in step (1) the corresponding cropped image will be extracted from the input image and fed to the second CNN. Since CNN models have fixed input sizes and the spots/eyespots have a wide range of sizes, all patches are resized to a common size before being fed to the measurements CNN. The CNN then performs the segmentation of the pattern element and outputs binary segmentation masks corresponding to each of the pattern elements we wish to measure.

In Fig 2, a basic scheme with the overview of the entire detection + measuring system is illustrated for an eyespot-bearing butterfly.

**Fig 2.**
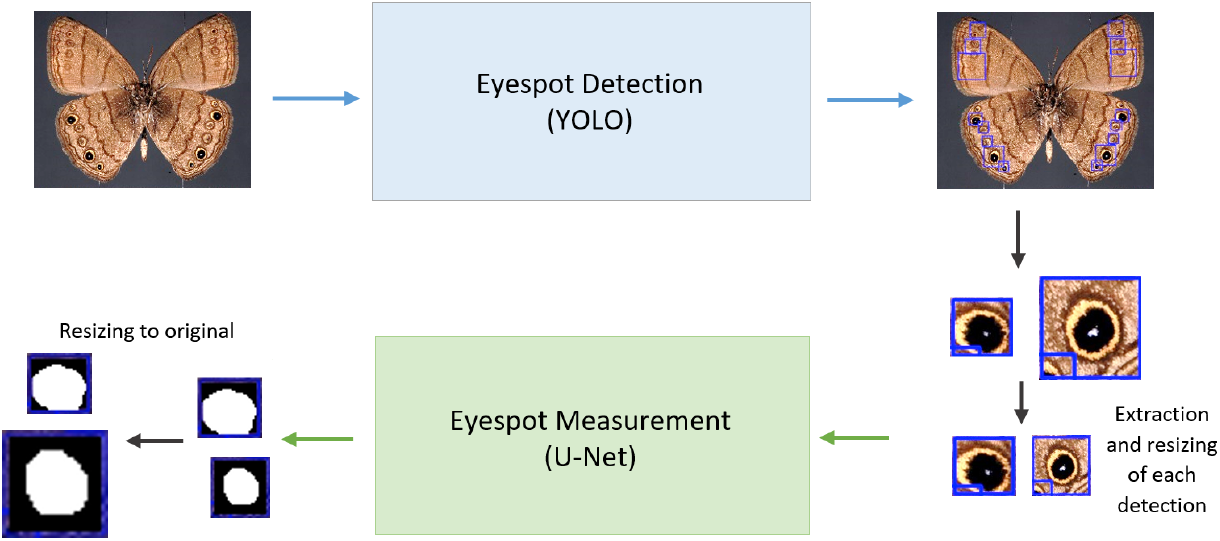
Overview of the two CNNs approach for first detecting and then measuring, spots or eyespots on images of butterfly wings.

#### Spot and eyespot detection

For spot and eyespot detection we used YOLO [23], a state of the art deep learning algorithm for object detection and classification. YOLO detects objects in the images and provides, for each detected object, a bounding box and class probabilities. YOLO is fast, compared to other object detection methods, because it uses the same CNN to predict the bounding boxes and the class probabilities for those boxes. Three alternatives were investigated, training YOLO with two classes, corresponding to the two types of pattern element, and training YOLO with only one class, either detecting all pattern elements without differentiating the type, or detecting only marginal eyespots, which are the most frequent, and the only ones present in dataset2.

#### Eyespot measurements

We treated the eyespot measurements problem as an image segmentation problem, and used a CNN to segment the different eyespot areas we wished to measure. Unlike previous approaches that only detect circular eyespots [14], this approach makes no assumptions on the shape of the eyespots.

For segmentation we used the U-Net [25], a widely used CNN designed for image segmentation. Although this encoder-decoder network was proposed, and is used primarily, for biomedical images, it can be used with images of any kind. Our U-Net was trained using patches containing the detected eyespots and not the whole image. The eyespot patches were resized, since U-Net inputs must be of fixed size. For each input patch, this network will output an image, of the same size, with binary segmentation masks for the different areas of the eyespot, namely the black and orange rings together, and the white center. Once an eyespot has been segmented, areas of the different segmentation masks are obtained by adding the pixels contained within the binary segmentation mask that is obtained at the output of the U-Net. It is also necessary to perform an inverse resize operation to the original dimensions that the crop of each eyespot had in its wing image. Once the area of the center and that of the surrounding color rings are determined, the total eyespot area is obtained by adding the two quantities.

We implemented two versions of the U-Net that can provide us with the same set of measurements, namely, area of the white center, and of the surrounding color rings. One segments only the two color rings from the rest of the image (two-class) and the other segments center, color rings, and background (three-class). Both versions are trained using the categorical cross entropy (CCE) as the cost function. We used the unweighted version of this cost function, which is more commonly used, and also a weighted version, termed Weighted Cross entropy (WCCE), which includes class weights. Using class weights is particularly important for the three-class model, where the center class has much fewer pixels, compared to the color rings and background classes. Using class weights in the loss function allows us to increase the weight of the center class to compensate for these differences.

The code for the YOLO CNN and the U-Net CNN is available for download at https://github.com/Margarida-Silveira/Butterfly_CNN.

#### Evaluation

YOLO models for the detection of the different pattern elements are evaluated using Average Precision (AP) and mean Average Precision (mAP). AP is the most commonly used metric to evaluate the performance of object detection algorithms. It is computed as the area under the Precision-Recall curve for Intersection over Union (IoU) thresholds between 0.5 and 0.95. mAP is the mean of AP for all the object classes, in this case the type of pattern elements (spots or eyespots).

Precision is the proportion of correctly classified pattern elements of a given type out of all the pattern elements identified by the model as that type, computed as:

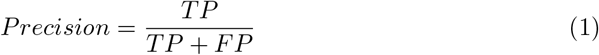

and Recall is the proportion of correctly classified pattern elements of a given type among all the pattern elements from that type, and is computed as:

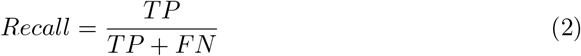

In the previous equation TP (true positives) is the number of correctly classified pattern elements from a given type and TN (true negatives) is the number of correctly classified samples from all the other types. Similarly, FP (false positives) is the number of incorrect classifications of a given pattern element type and FN (false negatives) corresponds to the total number of missed detections from all types of pattern elements.

A pattern element is considered a TP if the Intersection over Union (IoU) between its bounding box and any ground truth bounding box (manually identified) of the same type of pattern element is above or equal a given threshold, with IOU computed as the area of intersection divided by the area of union:

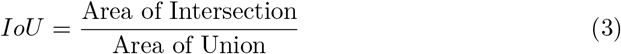

In case multiple detections of the same object occur, the one with highest probability is counted as a positive while the rest are counted as negatives.

U-Net segmentation results were evaluated with Accuracy and IoU. Accuracy measures the percentage of pixels that are classified correctly and IoU is computed in the same way as for YOLO detections, but in this case, it is computed between ground truth segmentation and predicted segmentation, and not bounding boxes. Finally, area measurements were obtained from the segmented images, converted to *mm*^2^ and compared to manual measurements.

## Results

### Eyespot detection

Dataset1 was randomly divided into 90% (3615 images) for training and 10% (402 images) for testing. For each image its pixel RGB values were divided by 255 to guarantee values in the 0 to 1 range. The results presented below were obtained by applying the YOLO trained models on the test data. This test set includes 402 images of whole butterflies with several spots/eyespots. In these butterfly images, there were 33953 pattern elements of the 2 types divided as shown in Table 2. Furthermore, 88% of all eyespots (14553 vs 2018) were marginal eyespots.

**Table 2.** Number of pattern elements of each type in the training/validation and test set for dataset1.

|  | Training Set | Test Set |
| --- | --- | --- |
| Spot | 15644 | 1738 |
| Eyespot | 14914 | 1657 |
| All | 30558 | 3395 |

Since only the right wing pattern elements were previously annotated, we manually annotated the elements in the left wing for the 402 test images using labellImg software [26]. To avoid the burden of also annotating the elements in the left wings for the training images, we instead erased the left part of each training image, as illustrated in Fig 3.

**Fig 3.**
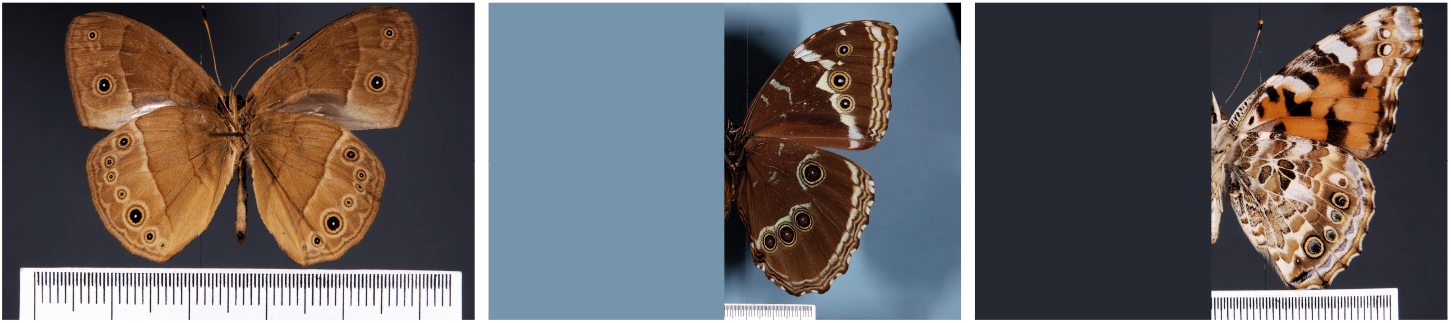
Examples of images from dataset1. Image of a complete butterfly and two butterfly images with left wing erased.

The YOLO network used was a publicly available implementation released in GitHub repository [24]. Fig 4 contains examples representative of the bounding boxes provided by YOLO models for the two pattern elements type classes (spot and eyespot), one class (where any of the types of pattern elements is detected), and one class marginal (where only marginal eyespots are detected).

**Fig 4.**
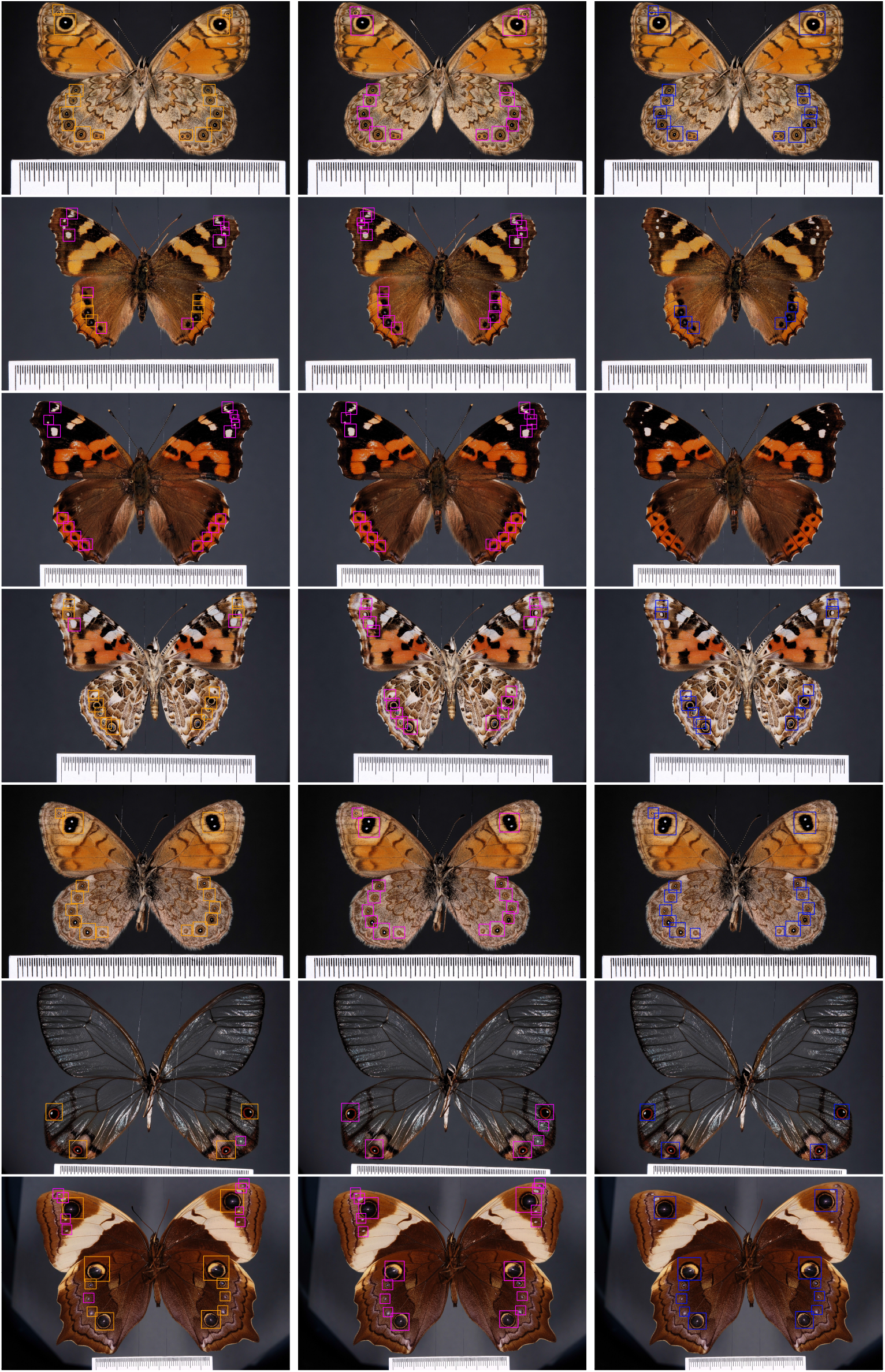
Detection examples. Left column shows detections of the two-class YOLO (spots in pink and eyespots in orange), middle column shows detections of one-class (no distinction between spot or eyespot types) YOLO and right column shows detections of “Marginal eyespots only” YOLO.

The network was trained with the Adam optimizer using an initial learning rate of 1e-04, *β*_1_ = 0.9 and *β*_2_ = 0.9999, and a batch size of 8. Training was stopped when the number of iterations reached 100 or earlier, if the loss function did not decrease for the validation set for more than 10 epochs (the patience hyperparameter). Duplicate detections were removed using Non Maximum Suppression (NMS) with a threshold of 0.45.

We trained YOLO using online data augmentation to increase the diversity of our training data. Our data augmentation strategy included horizontal flipping to make sure the network was trained with left-wing eyespots also and not affected by the removal, from the training images, of the wings that were not annotated.

The average precision (AP) and the mean average precision (mAP) values achieved in the test data are presented in Table 3.

**Table 3.**
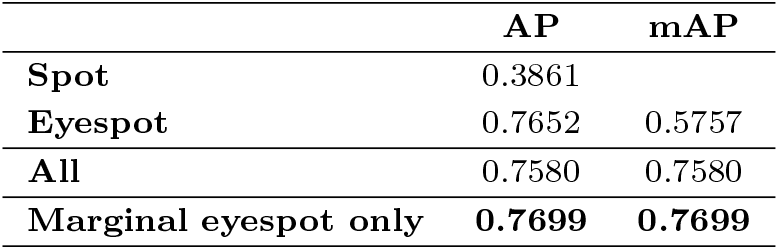
Scores achieved by the three YOLO models for eyespot detection on the test data from dataset1.

|  | AP | mAP |
| --- | --- | --- |
| Spot | 0.3861 |  |
| Eyespot | 0.7652 | 0.5757 |
| All | 0.7580 | 0.7580 |
| Marginal eyespot only | **0.7699** | **0.7699** |

We also tested the YOLO models trained with data from dataset1 on images from dataset2. The results are presented in Table 4.

**Table 4.**
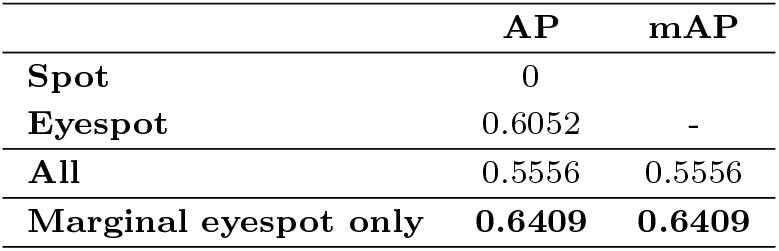
Scores achieved by the three YOLO models for eyespot detection on dataset2.

|  | AP | mAP |
| --- | --- | --- |
| Spot | 0 |  |
| Eyespot | 0.6052 | - |
| All | 0.5556 | 0.5556 |
| Marginal eyespot only | **0.6409** | **0.6409** |

### Eyespot measurements

To train the U-Net to perform automatic eyespot measurements we randomly divided dataset2 into 80% images for training (101 eyespots) and 20% images for testing (24 eyespots). We then used ground thruth area measurements from these 101 previously measured eyespots, for training. From the center X,Y coordinates and total area annotations in dataset2 we were able to create square eyespot cropped images, which were resized to 128×128 pixels. For each crop, we created ground truth binary segmentation masks with white circles corresponding to the eyespot color rings (black and orange rings combined) and to the center. The radius of each circle was obtained from the corresponding area manual measurements, after conversion from *mm*^2^ to pixels, and the position of the center was corrected manually. Fig 5 shows some examples.

**Fig 5.**
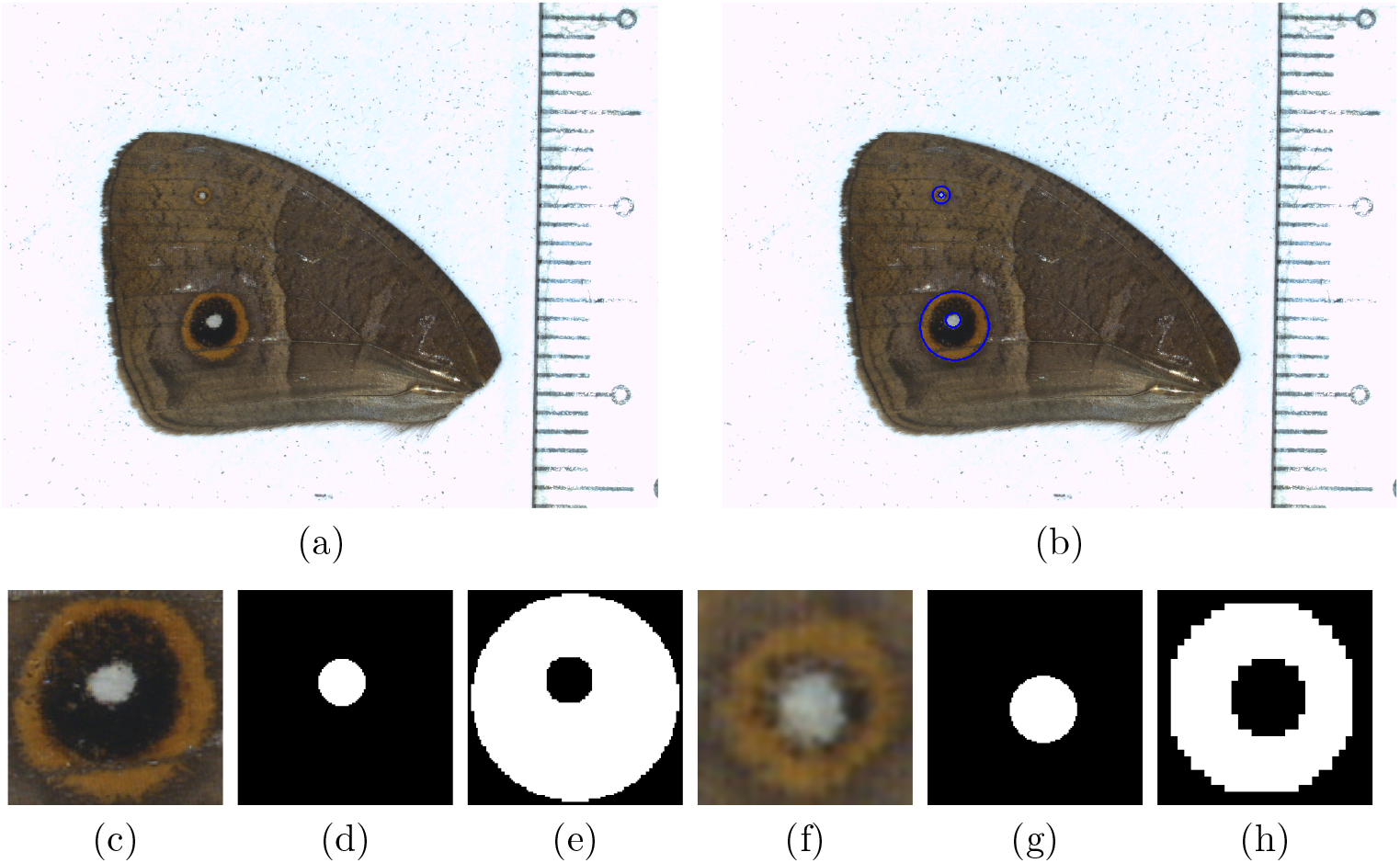
Illustration of ground truth created for *Bicyclus anynana* wing image, where white regions (bottom row) correspond to areas we wish to measure and for which we also have obtained manual measurements. (a) The original RGB image. (b) Original image with ground truth circles superimposed on each eyespot. (c) Large eyespot crop resized to 128×128 pixels. (d) Large eyespot center ground truth segmentation mask. (e) Large eyespot rings ground truth segmentation mask. (f) Small eyespot crop resized to 128×128 pixels. (g) Small eyespot center ground truth segmentation mask. (h) Small eyespot rings ground truth segmentation mask.

The results presented below were obtained by applying the U-Net trained models on the test set. The class weights used in WCCE loss function for two-class data were 1.75 and 1, corresponding to the eyespot rings area and to the rest of the image area, respectively, and for the three-class data they were 2, 25 and 1, for the eyespot rings, center, and background area classes, respectively. For the two-class model, to isolate the center region, we computed the negative of the segmentation mask and then eliminated the components connected to the image boundary using mathematical morphology.

Our U-Net was also trained with the Adam optimizer with an initial learning rate of 1e-04, *β*_1_ = 0.9 and *β*_2_ = 0.9999, and a batch size of 8. The network was trained for a maximum of 50 epochs, using an early stop criterion monitoring the validation loss with a patience of 10.

Table 5 presents the accuracy and IoU results of U-Net segmentation experiments with two-class data and with three-class data. In Fig. 6 we illustrate some of these results.

**Table 5.** Evaluation of U-Net segmentation models trained with different cost functions (CCE and WCCE). Acc = accuracy; F1 = F1-score; IoU = Intersection over union.

|  | Acc | F1 | IoU |
| --- | --- | --- | --- |
| Two-class CCE | 0.961 | 0.957 | 0.902 |
| Two-class WCCE | **0.963** | **0.960** | 0.912 |
| Three-class CCE | 0.953 | 0.914 | 0.908 |
| Three-class WCCE | 0.959 | 0.934 | **0.919** |

**Fig 6.**
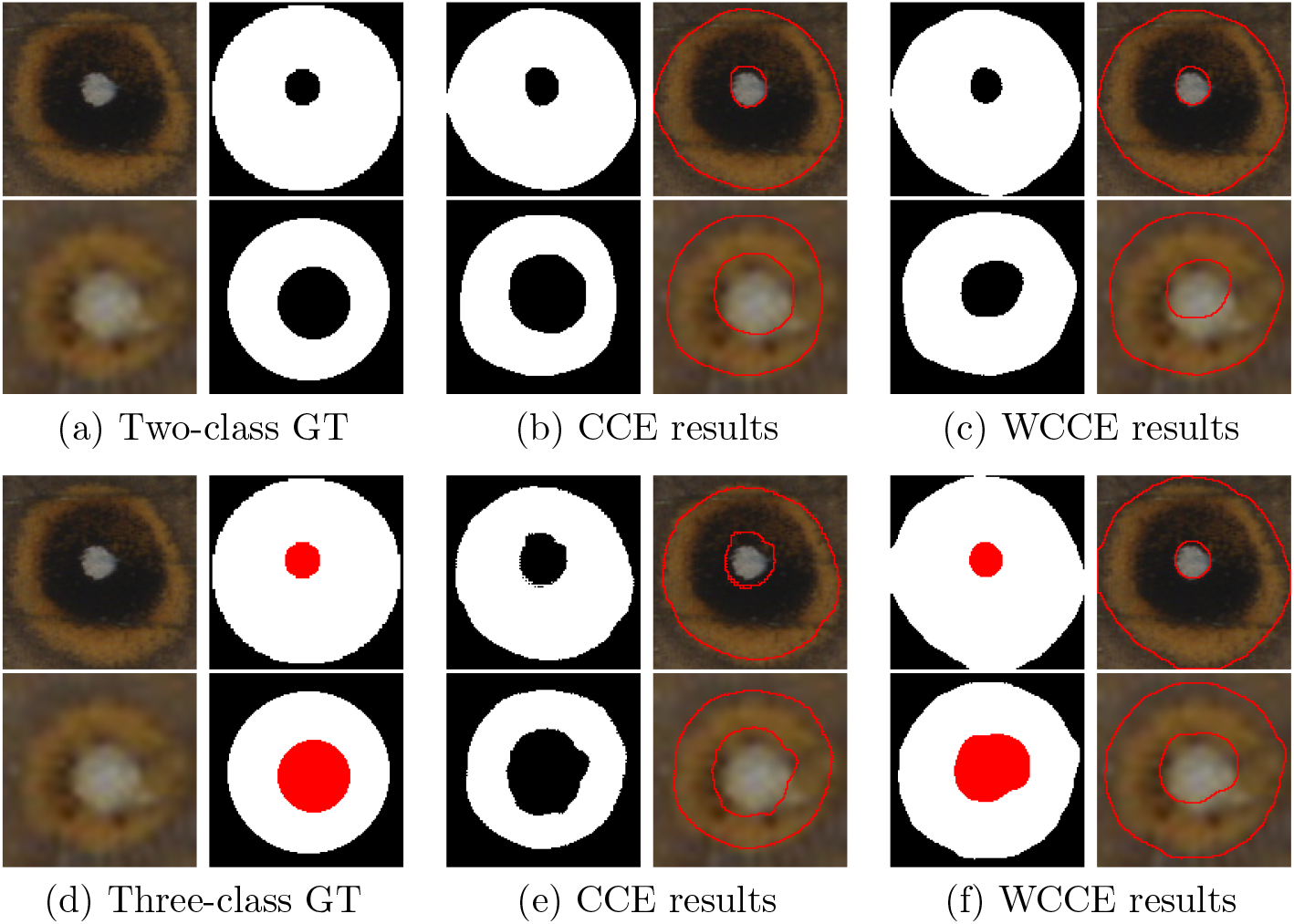
Example of U-Net segmentation results using different number of classes and cost functions. (a) original RGB eyespot image and its two-class ground truth segmentation, (b) predicted segmentation using CE loss function and corresponding contours. (c) predicted segmentation using weighted CE loss function and corresponding contours. (d) original RGB eyespot image and its ground truth segmentation, (e) predicted segmentation using CE loss function and corresponding contours. (f) predicted segmentation using weighted CE loss function and corresponding contours

For the best segmentation models, using two classes and three classes, we measured the areas of the eyespot rings and center in pixels and converted those measurements to squared millimeters (1 *mm* = 28.944 px). Then we computed the errors between manual and automatic area measures. These results are presented in Table 6.

**Table 6.** Average error and error standard deviation between manual and automatic measurements of the total eyespot and center areas. Values presented in pixels and *mm*^2^.

|  | U-Net two-class WCE |  | U-Net three-class WCE |  |
| --- | --- | --- | --- | --- |
|  | Center | Total | Center | Total |
| **Avg. Error [px]** | 13.13 | -93.85 | -9,70 | 12.81 |
| **Error std [px]** | 18.48 | 90.97 | 23.44 | 142.27 |
| **Avg. Error [$mm^2$]** | 0.02 | -0.11 | -0.01 | 0.02 |
| **Error std [$mm^2$]** | 0.02 | 0.11 | 0.03 | 0.17 |

## Discussion

YOLO was able to correctly detect a large number of spots and eyespots, despite the variability in the shape and color of these patterns and in the number of pattern elements per image. YOLO detected spots/eyespots even in cases where there is little contrast between these patterns and the background and in cases where spots/eyespots were overlapping. Most bounding boxes were very close to the ground truth ones, perfectly encapsulating the pattern elements, as illustrated in Fig 4. The majority of missed detections occurred for very small or overlapping spots/eyespots. Most of the false positives appeared in the area between two close eyespots. There were also a few detections that were counted as false positives but that corresponded to eyespots that were not included in the manually annotated ground truth. From the analysis of table 3 we can conclude that the two-class YOLO obtained better results for eyespots than for spots, probably because spots have higher pattern variability. With all data belonging to the same class, the YOLO network only has the task of detecting where the pattern elements are, without having to find out which class they belongs to. Thus, the mAP obtained with one class YOLO was 0.76, higher than the mAP obtained with two class YOLO of 0.58. The YOLO model trained to detect marginal eyespots achieved an AP of 0.77 outperforming the other two models tested here.

The results of the YOLO models to detect the eyespots in dataset2, shown in table 4 are not as good as the results of these models for the eyespots in dataset1, which was foreseable, since the training set (dataset1) has no examples from this butterfly species. The best model was “Marginal eyespots only”, as would be expected, since dataset2 only contains eyespots of this type. This model was able to achieve an AP of 0.64 on images from dataset2.

The U-Net models were able to segment the two areas within an eyespot, white center and black and orange rings combined, even though the training set was small and the ground truth was created assuming circular regions. Both models (two-class and three-class) had similar overall segmentation performance as shown in Table 5, achieving accuracies over 95%, F1-score over 91% and IoU over 90%. In the two-class model, using a weighted loss had little impact on the segmentation scores. In the three-class model, using the weighted categorical entropy was very important for accurately segmenting the small eyespot center, from the color rings and the background. The differences between manual area measurements and automatic ones are small, under 0.02*mm*^2^ for every model (both eyespots combined), as shown in Table 6. The best model for total area measurements was the weighted three-class approach but for center area measurements both methods performed equally well.

## Conclusion

In this work we investigated the use of convolutional neural networks to automatically detect and measure eyespots in images of butterflies. One CNN was trained to identify and distinguish spots and eyespots across the whole wing or marginal spots alone, in whole photos of different species of butterfly. Another CNN was trained to segment eyespot patterns into two different areas (center and remaining rings) in photos of a single species of satyrid butterfly. The accuracy of the identification and the precision of the measurements, when compared with that of manual identification and measurements was high. These CNNs, once implemented together with a graphical user interface, where imperfect pattern element detections can be manually corrected, can substantially accelerate the pace of research surrounding the ecology, evolution, and development of spots and eyespots in butterflies.

## References

1. Stevens M, Ruxton GD. Do animal eyespots really mimic eyes? Current Zoology. 2014, 60:26–36.

2. Kodandaramaiah U. The evolutionary significance of butterfly eyespots. Behav. Ecol. 2011, 22:1264–71.

3. Robertson K, Monteiro A. Female Bicyclus anynana butterflies choose males on the basis of their dorsal UV-reflective eyespot pupils. P Roy Soc Lond B Bio. 2005, 272:1541–6.

4. Stevens M. 2005. The role of eyespots as anti-predator mechanisms, principally demonstrated in the Lepidoptera. Biol. Rev. 2005, 80:1–16.

5. Oliver JC, Tong X-L, Gall LF, Piel WH et al. A single origin for nymphalid butterfly eyespots followed by widespread loss of associated gene expression. PLoS Genet. 2012, 8.

6. Oliver JC, Beaulieu JM, Gall LF, Piel WH et al. 2014. Nymphalid eyespot serial homologs originate as a few individualized modules. Proc. R. Soc. B. 2014, 281:pii: 20133262.

7. Huq M, Bhardwaj S, Monteiro A. Male Bicyclus anynana butterflies choose mates based on their ventral UV-reflective eyespot centers. J. Insect Sci. 2019, 19:25.

8. Prudic KL, Stoehr AM, Wasik BR, Monteiro A. Eyespots deflect predator attack increasing fitness and promoting the evolution of phenotypic plasticity. Proc. R. Soc. B. 2015, 282:20141531.

9. Bhardwaj S, Prudic KL, Bear A, Dasgupta M et al. Sex Differences in 20-Hydroxyecdysone Hormone Levels Control Sexual Dimorphism in Bicyclus anynana Wing Patterns. Mol. Biol. Evol. 2018, 35:465–72.

10. Monteiro A, Tong XL, Bear A, Liew SF et al. Differential Expression of Ecdysone Receptor Leads to Variation in Phenotypic Plasticity across Serial Homologs. PLoS Genet. 2015, 11.

11. Ozsu N, Chan QY, Chen B, Das Gupta M et al. Wingless is a positive regulator of eyespot color patterns in Bicyclus anynana butterflies. Dev. Biol. 2017, 429:177–85.

12. Matsuoka Y, Monteiro A. Hox genes are essential for the development of eyespots in Bicyclus anynana butterflies. Genetics. 2021, 217:1–9.

13. Conserved roles of Distal-less and spalt in regulating butterfly melanic patterns JLQ Wee, TD Banerjee, A Prakash, KS Seah, A Monteiro. bioRxiv. 2021.

14. Silveira, M., Monteiro, A. Automatic recognition and measurement of butterfly eyespot patterns. Biosystems. 2009.

15. Goodfellow, I.J.; Bengio, Y.; Courville, A., Deep Learning, MIT Press: Cambridge, MA, USA. 2016.

16. Jiuxiang Gu, Zhenhua Wang, Jason Kuen, Lianyang Ma, Amir Shahroudy, Bing Shuai, Ting Liu, Xingxing Wang, Gang Wang, Jianfei Cai, Tsuhan Chen. Recent advances in convolutional neural networks, Pattern Recognition. 2018, Volume 77.

17. Z. Li, F. Liu, W. Yang, S. Peng and J. Zhou. A Survey of Convolutional Neural Networks: Analysis, Applications, and Prospects. IEEE Transactions on Neural Networks and Learning Systems. 2021.

18. Kaya, Y., Kayci, L. Application of artificial neural network for automatic detection of butterfly species using color and texture features. Vis Compu. 2014, 30, 71–79

19. N. N. Kamaron Arzar, N. Sabri, N. F. Mohd Johari, A. Amilah Shari, M. R. Mohd Noordin and S. Ibrahim, Butterfly Species Identification Using Convolutional Neural Network (CNN). 2019 IEEE International Conference on Automatic Control and Intelligent Systems (I2CACIS), pp. 221–224.

20. F. Fauzi, A. E. Permanasari and N. Akhmad Setiawan. Butterfly Image Classification Using Convolutional Neural Network (CNN). 2021 3rd International Conference on Electronics Representation and Algorithm (ICERA), pp. 66–70.

21. Ayad Saad Almryad, Hakan Kutucu. Automatic identification for field butterflies by convolutional neural networks, Engineering Science and Technology, an International Journal. 2020, Volume 23, Issue 1, Pages 189–195.

22. Xin, Dongjun, Yen-Wei Chen, and Jianjun Li. Fine-Grained Butterfly Classification in Ecological Images Using Squeeze-And-Excitation and Spatial Attention Modules. Applied Sciences. 2020, no. 5: 1681.

23. Redmon, J., Farhadi, A., YOLOv3: An Incremental Improvement, arXiv 2018.

24. https://github.com/experiencor/keras-yolo3,

25. Ronneberger, O., Fischer, P., Brox, T. U-Net: Convolutional Networks for Biomedical Image Segmentation. MICCAI 2015.

26. Tzutalin. LabelImg. Git code. 2015. https://github.com/tzutalin/labelImg.

27. A. Voulodimos, N. Doulamis, A. Doulamis, and E. Protopapadakis. Deep learning for computer vision: A brief review, Computational intelligence and neuroscience. 2018.

